# AsaruSim: a single-cell and spatial RNA-Seq Nanopore long-reads simulation workflow

**DOI:** 10.1101/2024.09.20.613625

**Authors:** Ali Hamraoui, Laurent Jourdren, Morgane Thomas-Chollier

## Abstract

**Motivation:** The combination of long-read sequencing technologies like Oxford Nanopore with single-cell RNA sequencing (scRNAseq) assays enables the detailed exploration of transcriptomic complexity, including isoform detection and quantification, by capturing full-length cDNAs. However, challenges remain, including the lack of advanced simulation tools that can effectively mimic the unique complexities of scRNAseq long-read datasets. Such tools are essential for the evaluation and optimization of isoform detection methods dedicated to single-cell long read studies.

**Results:** We developed AsaruSim, a workflow that simulates synthetic single-cell long-read Nanopore datasets, closely mimicking real experimental data. AsaruSim employs a multi-step process that includes the creation of a synthetic UMI count matrix, generation of perfect reads, optional PCR amplification, introduction of sequencing errors, and comprehensive quality control reporting. Applied to a dataset of human peripheral blood mononuclear cells (PBMCs), AsaruSim accurately reproduced experimental read characteristics.

**Availability and implementation:** The source code and full documentation are available at: https://github.com/GenomiqueENS/AsaruSim.

**Data availability:** The 1,090 Human PBMCs count matrix and cell type annotation files are accessible on zenodo under DOI: 10.5281/zenodo.12731408.

## Introduction

Single-cell RNA sequencing technologies (scRNAseq) have revolutionized our understanding of cell biology, providing high-resolution insights on Eukaryote cellular heterogeneity. Still, studying the heterogeneity at the level of isoforms and structural variations is currently limited. Traditional short-read sequencing coupled with single-cell technologies (commonly droplet-based scRNA-seq protocols such as 10X Genomics) are not suitable for studying full-length cDNAs, because they require RNA/cDNA fragmentation, often resulting in the loss of information regarding the complete exonic structure (Arzalluz-Luque et Conesa 2018). Combining long-read sequencing, such as Oxford Nanopore or Pacbio, with single-cell technologies has enabled addressing this challenge (Arzalluz-Luque et Conesa 2018). Despite its advantages, the quality of Nanopore sequencing used to be impacted by higher error rates compared to short-read technologies, thus negatively impacting the detection of cell barcodes (CBs) and unique molecular identifiers (UMIs) (Karst et al. 2021). Yet, these elements are critical for attributing reads to their original cells, and for the accurate characterization and quantification of isoforms. That is why a hybrid approach, coupling long-read and short-read technologies, used to be necessary for a reliable assignment of CBs and UMIs (Lebrigand et al. 2020). Recently, the accuracy of Nanopore reads has been drastically improved (95-99% with the R10.3 flow cells (Dippenaar et al. 2021)), paving the way to untie long-read from short-read approaches in single-cell studies. Recently-released bioinformatics methods, including scNapBar (Wang et al. 2021), FLAMES (Tian et al. 2021), BLAZE (You et al. 2023), Sicelore 2.1 (Lebrigand et al. 2020), Sockeye (https://github.com/nanoporetech/sockeye) and scNanoGPS (Shiau et al. 2023), have been developed to detect CBs and/or UMIs without using companion short-read data (referred to as Nanopore-only methods). These advances have the potential to reduce both the cost and the amount of work traditionally associated with hybrid sequencing computational workflows.

In the context of these developments, evaluating Nanopore-only methods for processing single-cell long-read datasets remains challenging. Most of the methods currently available are benchmarked against short-read datasets; this approach is not devoid of biases and is therefore considered to be an imperfect gold standard (Ziegenhain et al. 2022; Sun et al. 2024). One solution lies in the use of simulated datasets, which can mimic real experimental outcomes without the same biases as empirical methods. Simulated data provide a known ground truth—true cell barcodes (CBs), true unique molecular identifiers (UMIs). This ground truth can be exploited by method developers in various ways, such as tuning method parameters, validating results, benchmarking novel tools against existing methods, and highlighting their performance across a wide range of scenarios. Besides, the focus of most long read scRNA-seq and spatial methods is to identify alternative splicing events and differentially expressed isoforms (DEI) between cell types or cell states (Joglekar et al. 2023). Assessing the performance of these methods is also challenging because the ground truth is typically not known, and simulating random reads without any biological insight does not address this issue. One solution to this issue is to use instead simulated datasets, in which the ground truth (e.g., DEI, Fold change, batch effect) is known.

To date, no existing workflow has been designed with the specific purpose of simulating single-cell or spatial RNAseq long-read data, especially with biological insights. A general workflow for long-read transcriptomic datasets, TKSM (Karaoğlanoğlu et al. 2024), comprises some modules that enable users to assemble a pipeline for scRNAseq, but it is not primarily intended for single-cell applications. Current scRNAseq counts simulation tools (such as SPARSim (Baruzzo, Patuzzi, et Di Camillo 2020) or ZINB-WaVE (Risso et al. 2018)) generate only a synthetic single cell count matrix. The bottleneck lies in the generation of simulated raw reads. It is notable that some studies on single-cell long-read methods, such as those described in Wang et al. 2021 and You et al. 2023, have employed simulated data. As part of these studies, individual tools (e.g., SLSim; https://github.com/youyupei/SLSim) have been developed to generate artificial template sequences with random cDNA, and simulators such as Badread (Wick 2019) or NanoSim (Yang et al. 2017) are employed to introduce sequencing errors based on a predefined error model. While such tools can effectively be used to benchmark the accuracy of CB assignment algorithms, it does not account for the complexities of estimating a realistic complete single-cell long-read dataset. Such complexities include PCR biases and artifacts, sparsity, variability, and heterogeneity—characteristics intrinsic to single-cell and spatial data. Comprehensive simulation would allow for broader and more precise benchmarking of the performance of single-cell long-read bioinformatics tools.

To address this gap, we have developed AsaruSim, a workflow that simulates single-cell long-read Nanopore data. This workflow aims to generate a gold standard dataset for the objective assessment and optimization of single-cell long-read methods. The development of such a simulator alleviates the bottleneck in generating diverse *in silico* datasets by leveraging parameters derived from real-world datasets. This capability enables the assessment of method performance across different scenarios and refines pre-processing and analysis methods for handling the unique complexities of long-read data at the single-cell level.

## Methods

AsaruSim mimics real data by first generating realistic UMI counts using SPARSSim (Baruzzo, Patuzzi, et Di Camillo 2020), and then simulating realistic Nanopore reads using Badread (Wick 2019). Five major steps are implemented (Figure 1).

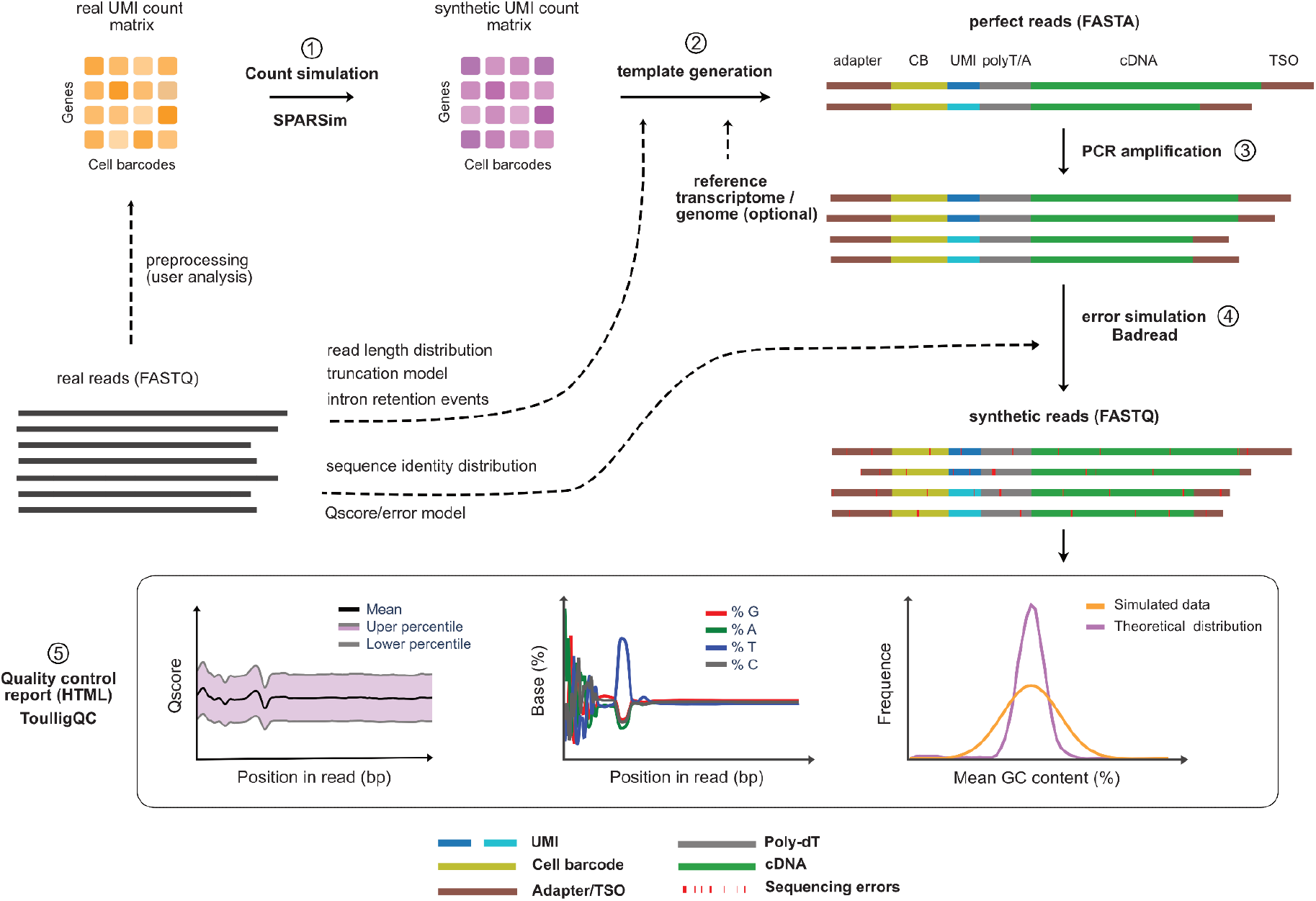

## 1 Synthetic UMI count matrix

AsaruSim takes as input a feature-by-cell (gene/cell or isoform/cell) UMI count matrix (.CSV), which may be derived from an existing single-cell short- or long-read preprocessed run, or from a count simulator tool. The R SPARSim library (Baruzzo, Patuzzi, and Di Camillo 2020) is used to estimate the count simulation parameters from the provided UMI count matrix and generate the corresponding synthetic count matrices, taking advantage of its ability to support various input parameters. AsaruSim also enables the user to input their own count simulation parameters, or alternatively, to select them from a predefined set of parameters stored in the SPARSim database.

### 2 Perfect raw reads generation

This step is an original Python script. AsaruSim generates synthetic reads based on the synthetic count matrix. The retro-engineering of reads is achieved by generating a corresponding number of random UMI sequences for each feature (gene or isoform). The final construction corresponds to a 10X Genomics coupled with Nanopore sequencing library (Lebrigand et al. 2020): an adaptor sequence composed of 10X and Nanopore adaptors, a cellular barcode (CB), UMI sequences at the same frequencies as in the synthetic count matrix, a 20-bp oligo(dT), the feature-corresponding cDNA sequence from the reference transcriptome, and a template switch oligo (TSO) at the end. When a gene expression matrix is provided, a realistic read length distribution is achieved by selecting a random transcript of the corresponding gene, with a prior probability in favor of short length cDNA (Supplementary Note a). An optional step can be performed to mimick unspliced reads by retaining introns (Supplementary Note b). In real data, reads are not always full-length as cDNA can be truncated. Here, each generated cDNA is thus truncated based on an empirically-derived truncation probability distribution, estimated by mapping a random subset of real reads to the reference transcriptome using Minimap2 (Supplementary Figure 6a,b), as described in Prjibelski et al. 2023. At the end, generated reads are randomly oriented, with each synthetic read having an equal probability of being oriented in the original strand or the reverse strand. These final sequences are named “perfect reads’’ as they exactly correspond to the introduced elements (CB, UMI, cDNA…) without addition of sequencing errors.

### 3 Mimicking PCR amplification bias (optional)

The perfect reads are duplicated through artificial multiple PCR cycles by an original Python script reimplemented from Sarkar, Srivastava, and Patro 2019; Orabi et al. 2019 with several optimizations to improve speed and memory usage. This enables us to take into account the bias of amplification introduced during library constructions Bolisetty, Rajadinakaran, and Graveley 2015. At each cycle, a synthetic read has a certain probability to be successfully replicated. The efficiency rate of duplication is fixed by the user (default p_dup_= 0.9). Then, each nucleotide in the duplicated read has a probability to be mutated during the process. The error rate is also fixed by the user (default p_error_*=* 3.5e-05). From this resulting artificial PCR product, a random subset of reads is finally selected to mimic the experimental protocol where only a subset of the sample is used for the sequencing step.

### 4 Introduction of sequencing errors in the reads

The perfect reads or post-PCR reads are used as template for Badread error simulation, which simulates Nanopore sequencing errors and assigns per-base quality scores based on pre-trained error models and sequence identity with the reference genome. AsaruSim allows the user to a) provide a personal pre-trained model, or b) providea real FASTQ read file to internally train a new model, or c) choose a pre-trained model within the Badread database. To approximate the observed sequence identity distribution in the experimental data, we align the real FASTQ read to the reference genome using Minimap2 (Li 2018), then calculate a sequence identity for each alignment from the Minimap2 output, with three possible identity models including or excluding gaps. A beta distribution is then fitted to the identity value to estimate the distribution parameters (Supplementary Note c).

### 5 Report

Finally, AsaruSim generates an HTML report presenting quality control plots obtained by analyzing the final FASTQ read files with ToulligQC (https://github.com/GenomiqueENS/toulligQC). This report aims to make sure the simulated data correspond to the expectations of the user before using them with tools dedicated to analyse scRNAseq long-read data.

AsaruSim is implemented in Nextflow (Di Tommaso et al. 2017) under GPL 3 license to allow a flexible and easily customizable workflow execution, computational reproducibility, and traceability (Supplementary Note d). To ensure numerical stability and easier installation, it also uses Docker (Merkel 2014) containerization technology.

## Results

We developed AsaruSim to produce artificial Nanopore scRNAseq data that resembles a real experiment in terms of biological insights.

As a use case, we used a public dataset of human peripheral blood mononuclear cells (PBMCs) (10X, 2022) as reference data. We downloaded the count matrix and used it as input to AsaruSim. From the 5,000 cells initially present in the original matrix, we selected 3 cell types (CD8+T, CD4^+^T and B cells) resulting in 1,090 cells then used as a template to simulate the synthetic UMI count matrix (step1). Next, we simulated 20 million perfect reads (FASTA) (step2) with 10 PCR cycles (step 3). We downloaded a subset of one million original FASTQ raw reads to generate the error model for Badread, and then introduced errors to generate the synthetic reads (FASTQ) (step4). The quality control report is finally generated (step5, Supplementary Note e).

We compared the properties of the simulated data to the experimental data. Both datasets showed similar a) read-length distribution and transcript coverage, b) number of mismatches and insertions/deletions in reads aligned to the 10X adapter sequence using VSEARCH (T et al. 2016)(Supplementary Note e).

Next, we pre-processed the simulated raw reads using the Sockeye pipeline (https://github.com/nanoporetech/sockeye), and both experimental and simulated matrices were processed using Seurat v5 (Hao et al. 2024). The correlation of the average log fold change for cell types markers between real and simulated data shows a pearson’s correlation coefficient r=0.84 and the integration of both datasets shows a miLISI=1.6, demonstrating a good agreement in gene expression between the real and simulated datasets (Supplementary Note e).

When comparing with TKSM (Karaoğlanoğlu et al. 2024), AsaruSim outperforms TKSM in terms of features specific to single-cell applications, similarity between real and simulated data, and computing efficiency (Supplementary Note f).

## Conclusion

We presented a comprehensive workflow for simulating single-cell Nanopore data from the matrix to the sequence level, to create custom gold standard datasets. Potential applications include generating reads with differential gene expression (DEG) or differential expression of isoforms (DEI) between cell groups, as well as simulating known fold changes or batch effects, to assess and optimize single-cell long-read methods. AsaruSim offers a variety of configuration options to allow for flexible input and design.

Currently, AsaruSim generates data compatible with the 10X Genomics 3’ protocol. We plan to expand AsaruSim to accommodate additional single-cell techniques and protocols, and support for PacBio sequencing.

## Author contributions

AH and MTC contributed to the design and implementation of the research, to the analysis of the results and to the drafting of the manuscript. LJ participated in testing the tool, accessibility in GitHub and documentation. All authors contributed to the final version of the manuscript.

## Acknowledgements

We thank Alice Lebreton for insightful discussions regarding this work.

## Supplementary Notes

### a Mimicking the real read length distribution (for use in step 2)

AsaruSim generates reads that correspond to a 10X Genomics coupled with Nanopore library construction (Figure 1). Regarding the cDNA sequences, the length of the reference annotation sequences may not mimic real datasets. Indeed, in the publicly available data generated with 10X Genomics devices coupled with Nanopore sequencing, it has been observed that the average read length does not exceed 1.2 kb (i.e. Tian et al. 2021; Shiau et al. 2023). Yet, many organisms (such as human or mouse) have an average cDNA length that exceeds this value; using the full length of the reference cDNA for the simulated data would thus not be realistic (Supplementary figure S1a).

If the user provides a gene expression matrix, a realistic read length distribution is achieved by selecting a random transcript of the corresponding gene, with a prior probability in favor of short length cDNA. To approximate the observed distribution of read lengths, we first identified the statistical distribution that best describes the real dataset. A number of candidate distributions (Weibull, gamma, log-norma) are fitted with the real data. Then, we calculated an Akaike information criteria AIC; *AIC = 2K - 2ln(L*), where K is the number of reads and L is the log-likelihood estimated from the probability density function. Based on the comparison of the AIC values, we determined that the log-normal distribution best approximates the read length distribution (Supplementary figure S1b).

This log-normal distribution is then used to efficiently approximate the cDNA length distribution of the real dataset. We implemented a model-fitting approach so that users may fit their data by providing a subset of real reads (in FASTQ format, Figure 1). Then, the perfect reads (step 2) are generated by selecting a random transcript of the corresponding gene. When a gene-by-cell matrix is provided, the choice of this reference cDNA is achieved by using the estimated log-normal distribution as a probability function to give more chance to transcripts having the length that matches the real read length distribution (thus favoring short length transcripts).

An alternative input to the feature-by-cell UMI count matrix is the submission of a per barcode UMI count CSV file to AsaruSim. The user thus only controls the cell number and sequencing depth of synthetic data. In this scenario, the genes will be randomly selected from the reference transcriptome following the estimated read length probability function.

**Figure S1.**
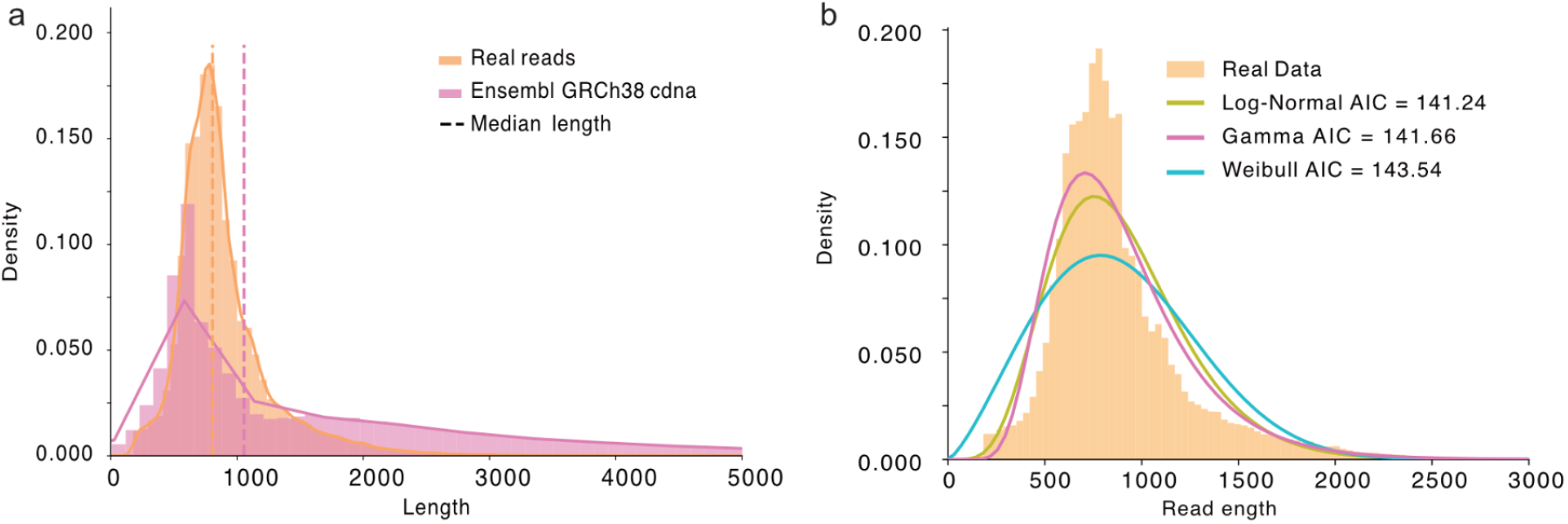
Read length distributions. **a**. Comparison between Ensembl cDNA (median 1055 pb) and real reads lengths (median = 806 bp). The real reads comprise the real cDNA sequence (median ~700 bp) and ~100 bp of various adaptor sequences. **b**. Fitting of theoretical distributions to real data. Different theoretical candidate distributions (Log-Normal, Gamma and Weibull) are fitted to the real read length distribution. The Akaike information criterion (AIC) shows that the Log-Normal distribution best fits the real read distribution.

### b Simulating unspliced reads and Intron retention events

Real single-cell, and especially single-nuclei, libraries contain a significant proportion of intronic reads—up to 20% in single-cell 3’ human PBMC datasets (10x Genomics, Technical Note CG000376). These reads can originate from unprocessed pre-mRNA and intron-retained transcripts, although the latter typically occur at lower proportions. To simulate intron retention events, we use a Markov chain model to represent the transitional probabilities between spliced and retained intron states, based on the state of the preceding intron, as described in NanoSim (Hafezqorani et al., 2020). This model is generated by running the model_intron_retention.py module from NanoSim on real FASTQ reads. Additionally, to simulate unprocessed pre-mRNA, AsaruSim allows users to specify a fraction of reads that will remain unspliced; in this fraction, all introns within the transcript are retained. This feature is demonstrated in Supplementary note e.

### c Sequence identity estimation (for use in step 4)

Badread uses the beta distribution to model the sequence identity distribution. In order to identify the optimal parameters (mean, standard deviation, and maximum) that best describe this beta distribution for our real data, a step search method was conducted throughout simulations of multiple subsets, each containing 100,000 reads simulated from the human reference transcriptome hg38. Briefly, in each step we compare the subset to the real data and rerun the simulation with one of the three parameters adjusted. To compare the subset to the real data, the 10X adapter sequence was aligned to the real and simulated reads using VSEARCH (T et al. 2016). The resulting sequence identities were then used to compute an MSE between the real and simulated reads:

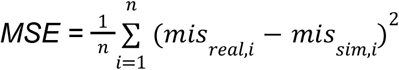

as *mis*_*real*,*i*_ and *mis*_*sim*,*i*_ are the number of mismatches from real and simulated data. Further details of this simulation can be found in our GitHub repository (https://github.com/GenomiqueENS/AsaruSim).

Sequence identity can be computed in multiple ways from a sequence alignment. We provide three identity models : the most commonly used method is “Gap-Excluded Identity”, which excludes all gapped positions from the alignment, calculated as:

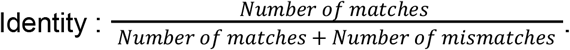

Alternatively, the “BLAST Identity” (Altschul et al., 1990) is defined as the number of matching bases divided by the total number of alignment positions. “Gap-Compressed Identity” treats consecutive gaps as a single difference. Even though Badread uses BLAST definition to define sequence identity, our simulation shows that using the “Gap-Excluded Identity” in our model best approximates the real sequence identity (Supplementary figure S2). That’s why we set as default the Gap-Excluded Identity model.

**Figure S2.**
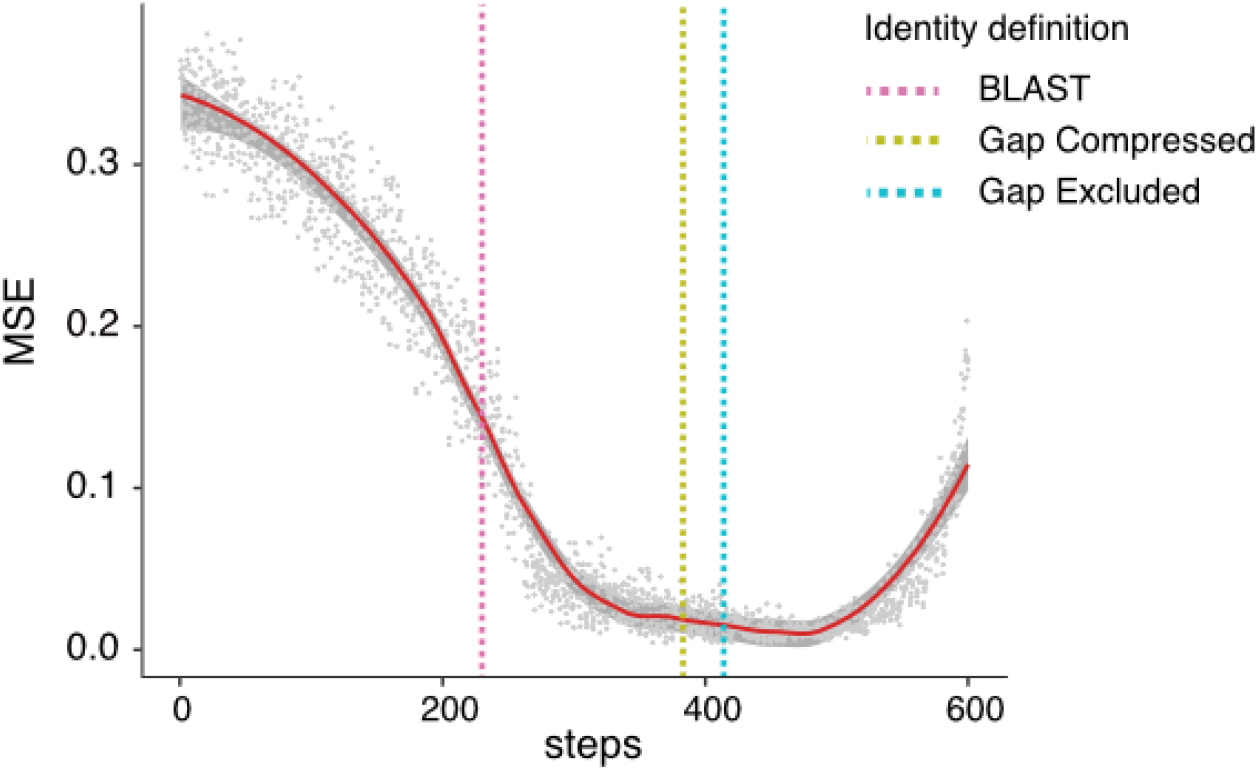
Parameter selection. Variation of mean square error (MSE) for sequence identity between real and simulated reads across different beta distribution parameters. To determine the optimal beta parameters that best approximate the sequence identity of real data, we conducted step search study through multiple simulations, adjusting one of the three parameters (mean, std, max) at each step. To measure the Mean Square Error (MSE) between real and simulated reads, we first aligned the adapter sequence to the real and simulated reads using VSEARCH (T et al. 2016). Then we used the resulting number of mismatches to calculate an MSE. Next, we plotted the parameters found with our fitted model using each definition of sequence identity (BLAST, Gap-compressed and Gap-excluded; vertical lines).

### d Nextflow implementation

AsaruSim is implemented in Nextflow to allow an easier installation and a robust processing and reproducibility. It allows flexible inputs; users may provide a UMI count matrix (.CSV), or a list of barcode counts (.CSV) or selection of the existing dataset from the SPARSim database. Additionally, users have the option to supply a reference raw read (.FASTQ) to estimate the error profile or choose from a wide range of predefined parameters. AsaruSim generates the simulated reads in FASTQ format and provides a comprehensive quality control report in HTML format (Supplementary Fig. S3).

**Figure S3.**
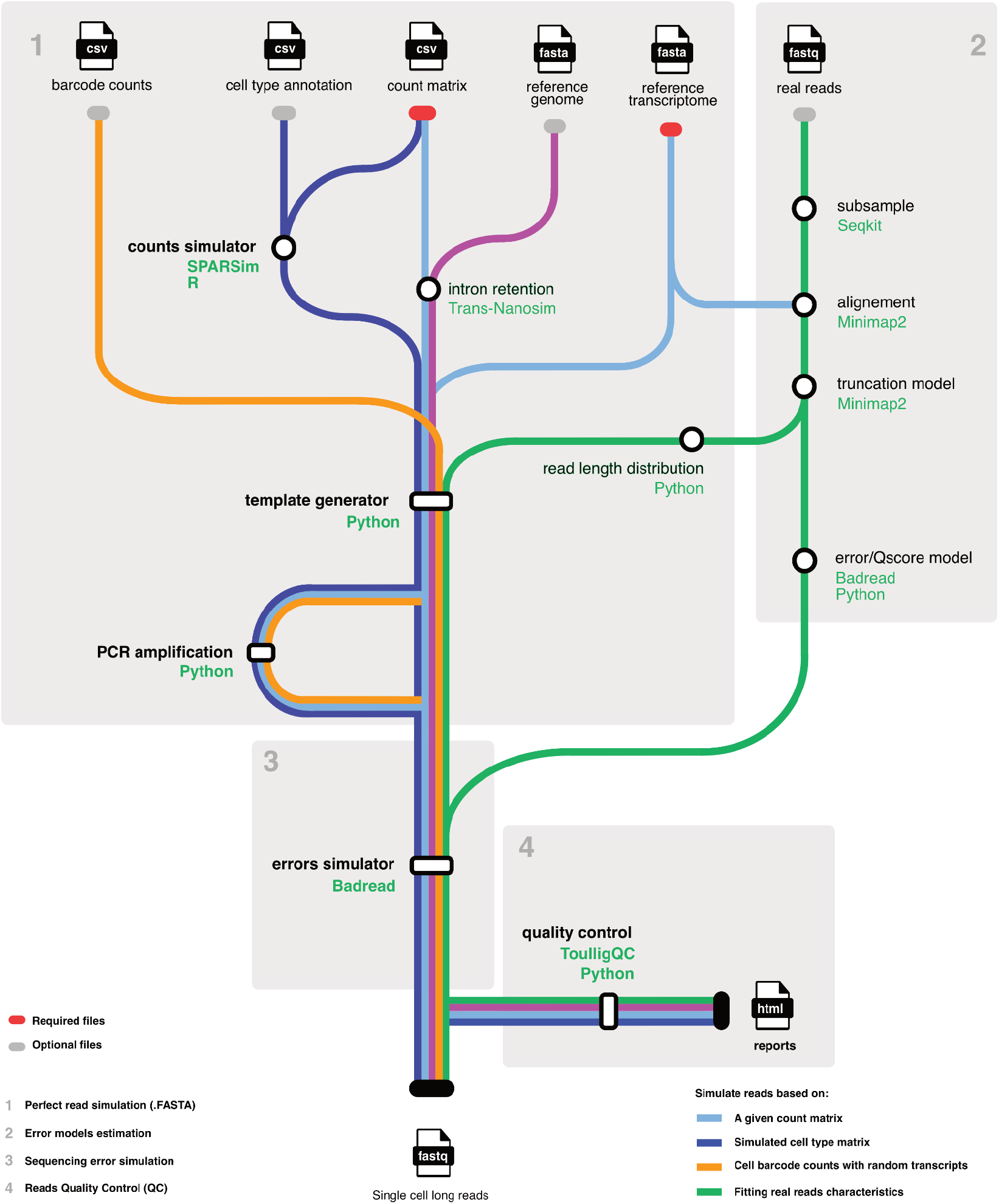
Workflow schema. Schematic representation of AsaruSim nextflow workflow showing the main 4 steps: 1-synthetic UMI count matrix, 2-perfect raw reads generation, 3-mimicking PCR amplification bias (optional), 4-introduction of sequencing errors in the reads, and 5-Quality control report.

### e Comparison of real *vs* simulated reads in the PBMC study case

In this section, we compared the characteristics of our simulated data (Supplementary Figure S4) with the real single cell long read data used as reference. The PBMC dataset, jointly produced by 10X Genomics and Nanopore was downloaded from (« 5k Human PBMCs, 3’ v3.1, Chromium Controller », s. d.).

**Figure S4.**
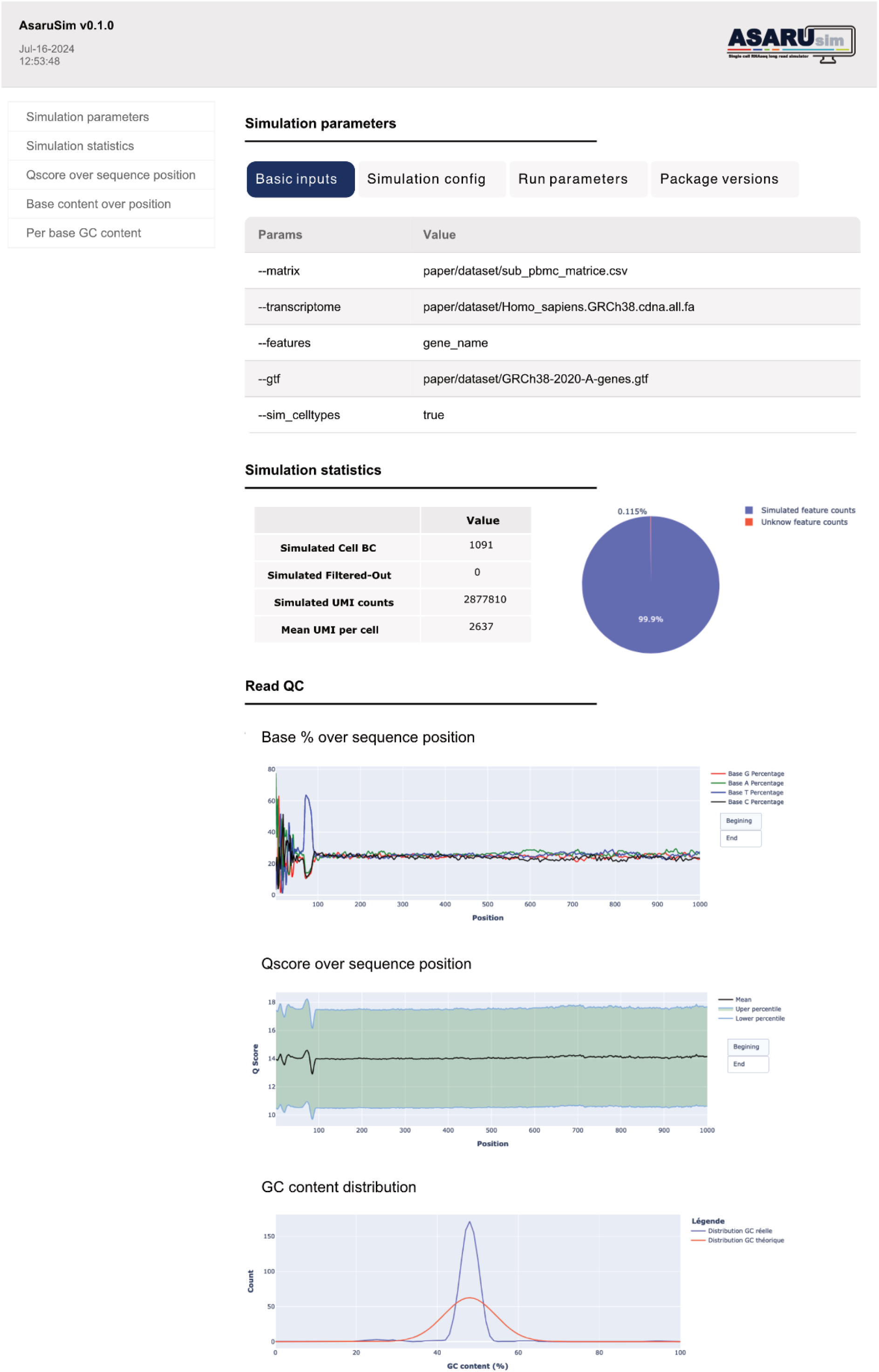
Quality control report: A capture of the HTML report generated by AsaruSim. The report is divided into three sections: Simulation parameters section, includes files and parameters used for the simulation to ensure traceability. Simulation statistics section, provides simulation statistics, including the number of BCs and UMIs, as well as general statistics about the run. Read QC section, provides the median Qscores, the percentage of bases over read sequences, and the GC content percentage.

We first compared our simulated data with real data in terms of read length distribution, transcript coverage, number of mismatch and number of substitutions (Supplementary Figure S5). We observed similar properties between the simulated and real datasets. The distribution of read lengths was similar across the two datasets (Supplementary Figure S5a). After alignment to the human hg38 genome, both datasets show comparable gene body coverage (Supplementary Figure S5b); the post-alignment QC results using Picard toolkit (Broad Institute., 2019) on simulated data combining intron retention events and 7% of random unspliced transcripts, indicate similar proportions of bases aligning to intronic, UTR, and coding regions (Supplementary Figure S5c).

**Figure S5.**
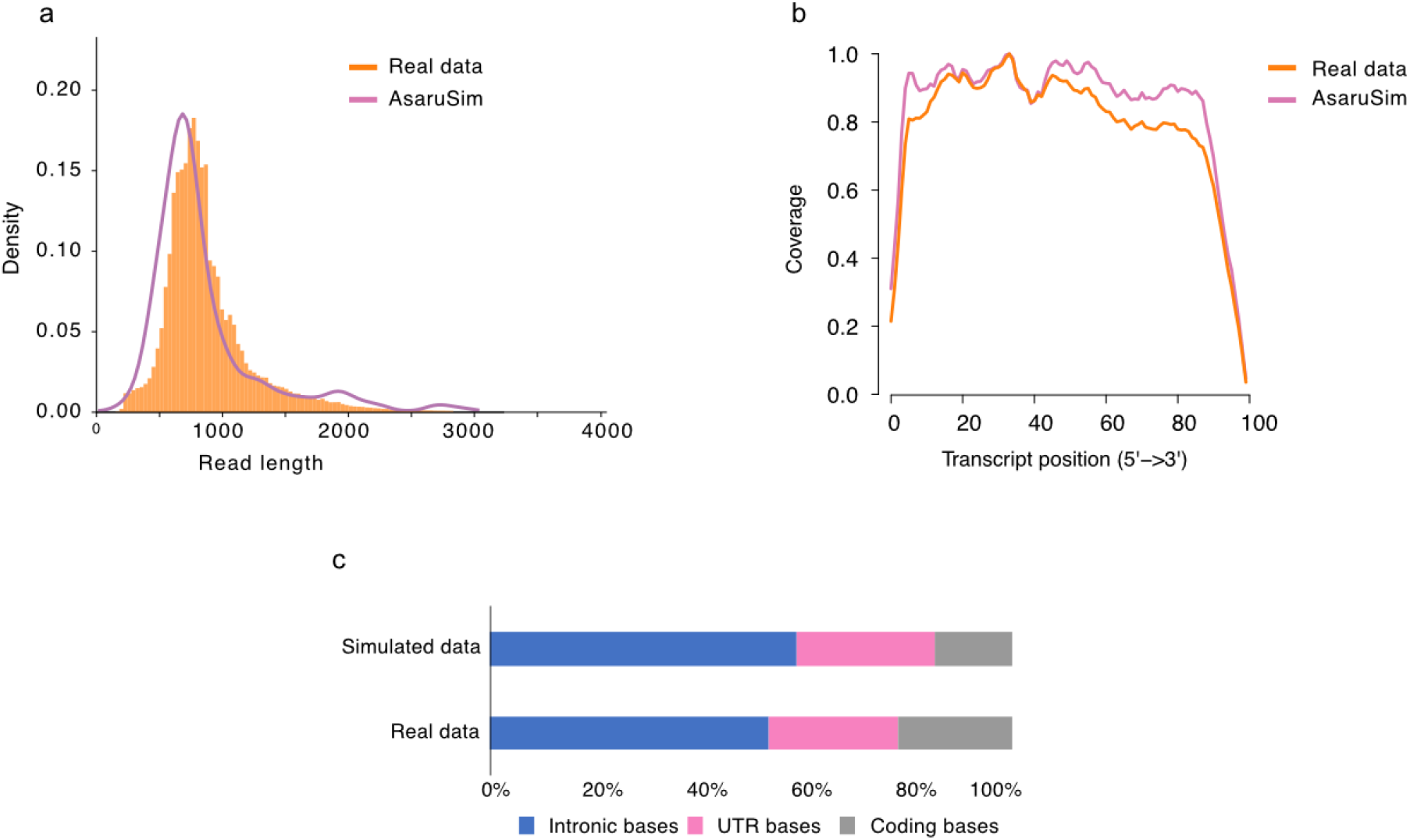
Comparison of real vs simulated reads. **a**. Read length distribution of the real and simulated reads. The distribution used for simulated cDNA size is obtained by fitting the read size distribution observed in the experimental data. **b**. Transcript body coverage plot for the simulated reads and real reads. This plot was generated with the Qualimap tool (Okonechnikov, Conesa, et García-Alcalde 2016) using as input the BAM file tagged by Sockeye workflow (nanoporetech, 2023). **c**. Post-alignment QC results compare the alignment of real and simulated datasets (combining intron retention events and 7% of random unspliced transcripts) to the human hg38 reference genome. Figure representing percent of bases aligned to intronic, UTR and coding regions.

After alignment to the 10X Genomics adapter sequence using VSEARCH (T et al. 2016)., the number of mismatches, indel errors and percentage of alignment identity (97%) revealed very similar between the real and simulated data (Supplementary Figure S6c, S6d). These results demonstrate the capability of AsaruSim to effectively simulate data that closely emulates real data.

**Figure S6.**
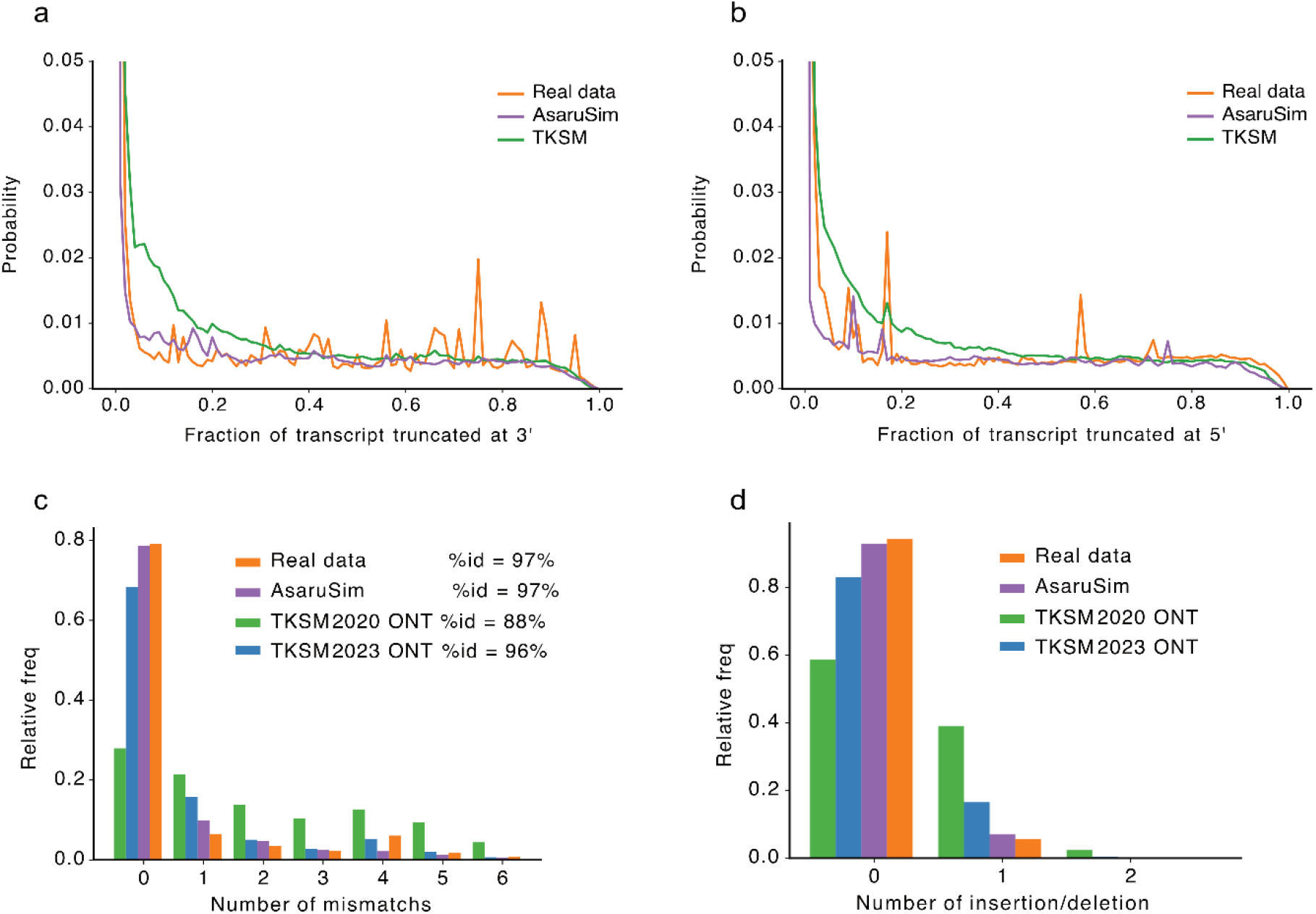
Comparison of AsaruSim vs TKSM simulated reads. **a**. Empirical 3’ truncation probability distributions for real human PBMCs ONT data (10X, 2022) versus data simulated using either AsaruSim or TKSM. The distributions were estimated by mapping reads to the human hg38 reference transcriptome using Minimap2. **b**. Same as panel a, but for the empirical 5’ truncation probability. **c**. Number of mismatches and percentage of alignment identity in reads aligned to 10X Genomics adapter sequence using VSEARCH (T et al. 2016), Two of Badread’s recommended read identity parameters were tested with TKSM: the default parameters (for nanopore R9.4.1 chemistry) and the latest model (95,99,2.5), corresponding to the R10.4.1 chemistry. **d**. Number of gaps in reads aligned to 10X Genomics adapter sequence using VSEARCH.

We then preprocessed the simulated raw reads and the real reads in a similar way using the Sockeye workflow (nanoporetech, 2023), and performed downstream analyses of the resulting gene count matrices with seurat V5. We compared the t-SNE visualisation of simulated and real data (Supplementary Figure S7a and S7c). miLISI measures the degree of mixing among datasets; the value of 1.6 shows a good closeness between the real and simulated data. Finally, we found a high correlation of the average log fold change for the cell types markers between real and simulated data (Pearson’s r=0.84). We conclude that our simulation workflow is robust to simulate single-cell RNA-seq long reads with gene expression profiles closely resembling those of real datasets.

**Figure S7.**
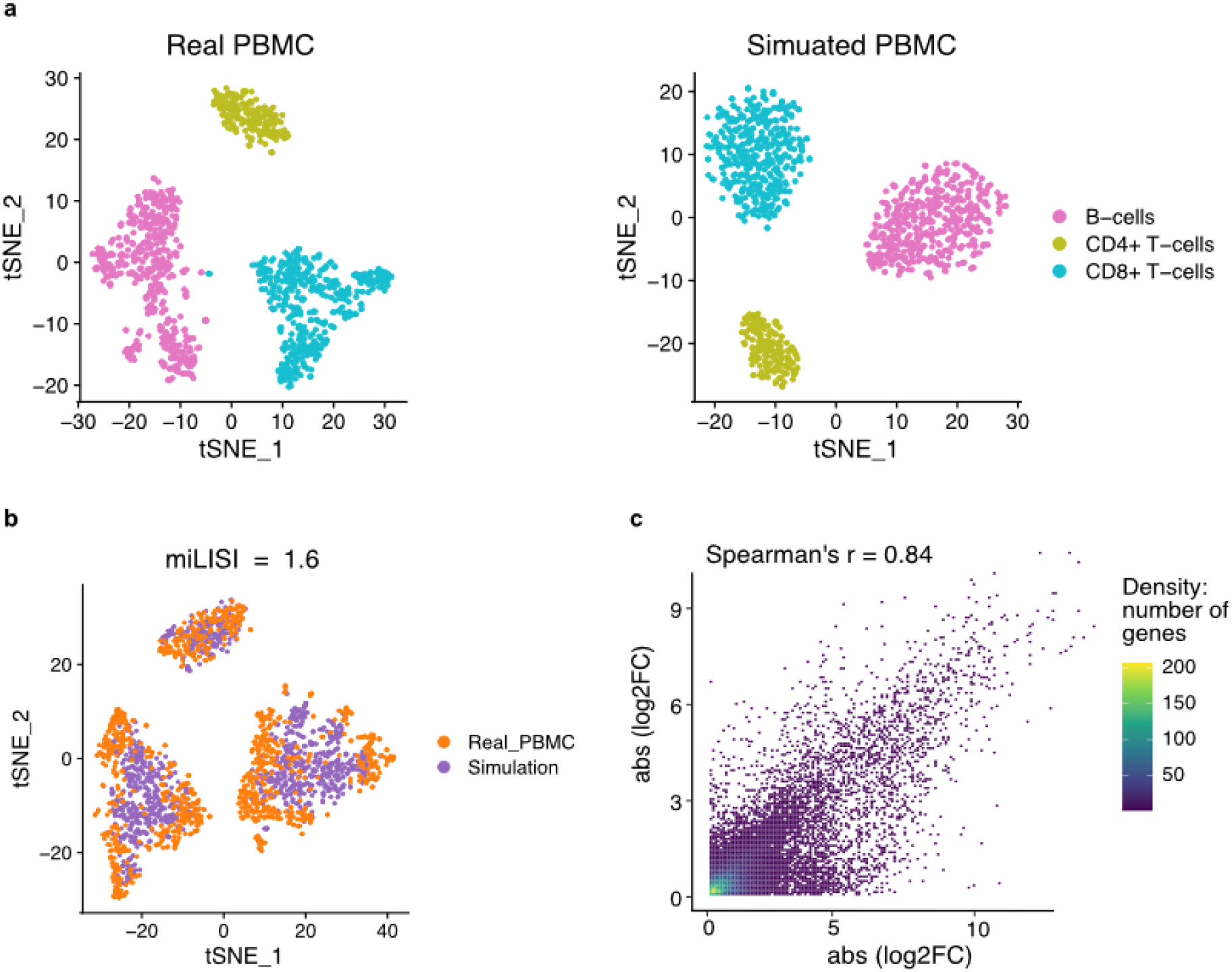
The t-SNE plots of real vs simulated gene expression count matrices. **a**. t-SNE representation based on the dataset of real PBMC cells (left) or AsaruSim simulated data (right). The matrices were processed using Seurat v5. For quality control purposes, genes expressed in less than three cells were discarded, and cells with gene counts more than 3,000 or fewer than 200 were filtered out. Then, cells were colored following the cell types annotation which was obtained using the singleR R package (Aran et al. 2019). **b**. Pairwise integration of two single cell datasets. The Seurat CCAIntegration algorithm was used for integration and the batch effect is removed using Seurat V5. The mean integration Local Inverse Simpson Index (miLISI) score was calculated using the LISI R package (Korsunsky et al. 2019). **c**. Density scatter plot shows the correlation of the absolute value of average log fold change (log2FC) for the cell types markers between real and simulated data. The log2FC values are computed using Seurat v5 FindMarker function.

### f Comparison between AsaruSim and TKSM

TKSM (Karaoğlanoğlu et al. 2024) is a transcriptomic sequencing long-read simulator providing modules that users can assemble into a pipeline, according to the sequencing design of the real dataset. Both AsaruSim and TKSM benefit from a workflow manager (Supplementary Table S1). In terms of features, both AsaruSim and TKSM take into account PCR cycles, Error and Qscore modeling, and truncation of reads modeling. TKSM supports gene/isoform fusion. In contrast to TKSM, AsaruSim provides unique features including read identity modeling, unspliced and novel transcripts simulation, as well as an HTML report to control the quality of the simulated dataset. Contrary to TKSM, AsaruSim does not take as input a FASTQ file of real reads, which users may not have, especially for public datasets that are available as already-processed count matrices. That’s why AsaruSim takes as input a count matrix (gene or isoform). A distinct feature of AsaruSim is its capability of simulating counts (step 1) with user’s personal simulation parameters, based on SPARSim (Baruzzo, Patuzzi, et Di Camillo 2020). For sake of comparison, Table S1 includes SLSim (You et al. 2023), which simulates basic reads, including only the Error and Qscore modeling.

**Table S1.** Comparison of AsaruSim versus TKSM and SLSim features. We have adapted TKSM to build a pipeline customized to the sequencing construction used to produce the PBMC study dataset (available on our reproduction package: https://github.com/alihamraoui/AsaruSim_Application_Note). We then simulated data with AsaruSim and TKSM, providing as input the FASTQ reads to TKSM and the count matrix to AsaruSim. We compared features common to both tools: Error and Qscore modeling as well as the truncation modelling (Supplementary Figure S6). The truncation profile of AsaruSim is more similar to the truncation profile of the real dataset than TKSM’s profile, in particular for the truncation probability distribution values, where TKSM tends to over-truncate read sequences. This observation applies for both 3’ and 5’ truncation (Supplementary Figure S6a,b). Regarding the error modelling, we aligned the reads to the 10X Genomics adapter sequence to calculate the number of mismatches, indel errors and percentage of alignment identity (Supplementary Figure S6c,d). The results obtained with AsaruSim are consistently more similar to the real data than TKSM, even when modifying TKSM to incorporate Badread’s read identity parameters for the latest R10.4.1 chemistry. These results demonstrate the higher capability of AsaruSim to simulate data that more accurately emulates real sequencing data.

|  | AsaruSim | TKSM | SLSim |
| --- | --- | --- | --- |
| Programming language | Python/R | C++/Python | Python |
| Workflow manager | Nextflow | Snakemake | **X** |
| Input | Gene/isoform counts matrix<br>CB counts<br>SPARSim presets | FASTQ | CB counts |
| Counts simulation | **V** | **X** | **X** |
| PCR simulation | **V** | **V** | **X** |
| Error and Qscore modeling | **V** | **V** | **V** |
| Read Identity modeling | **V** | **X** | **X** |
| Truncation modeling | **V** | **V** | **X** |
| Intron retention / unspliced transcripts | **V** | **X** | **X** |
| Gene/isoform fusion | **X** | **V** | **X** |
| Novel isoforms | **V** | **X** | **X** |
| Simulation QC report | **V** | **X** | **X** |

Last, we compared the computing efficiency of both tools (Supplementary Table S2). The runtime of AsaruSim is significantly lower than TKSM, thanks to a more efficient parallelisation of Badread. The peak memory is also lower for AsaruSim.

We compared the computational efficiency of both tools (Supplementary Table S2). AsaruSim demonstrates significantly lower runtime compared to TKSM, attributed to its more efficient parallelization of Badread. Additionally, AsaruSim requires less peak memory, further highlighting its computational advantages.

**Table S2.**
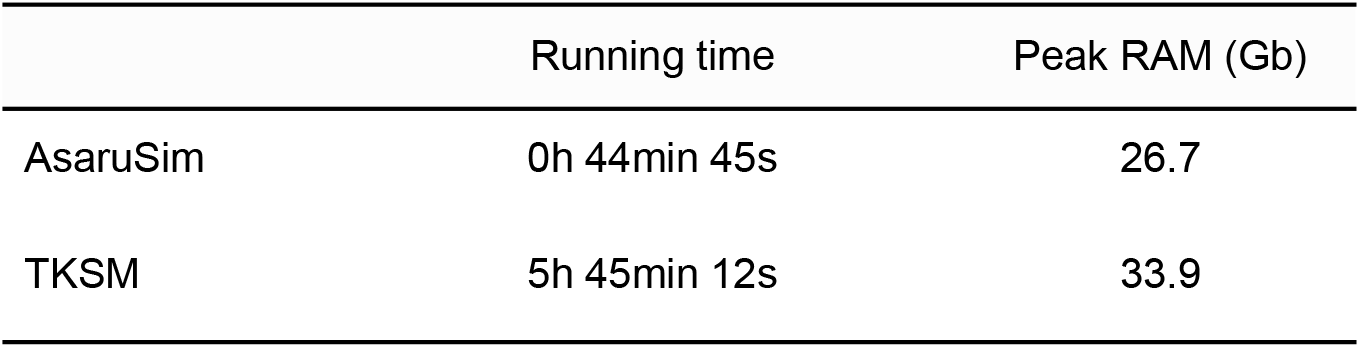
Comparison of AsaruSim and TKSM runtime and peak memory usage. Runtime and peak memory were measured using the Linux time command with the -v option during the simulation of 5,000 cells with 5 PCR cycles, totaling 5 million reads. Simulations were executed on the same machine (32 cores, 189 GB RAM) using 20 threads.

|  | Running time | Peak RAM (Gb) |
| --- | --- | --- |
| AsaruSim | 0h 44min 45s | 26.7 |
| TKSM | 5h 45min 12s | 33.9 |

